# Deciphering Spatially Resolved Pathway Heterogeneity in Ovarian Cancer Post-Neoadjuvant Chemotherapy

**DOI:** 10.1101/2025.08.08.669449

**Authors:** Akansha Srivastava, P K Vinod

## Abstract

High-grade serous ovarian cancer (HGSOC) is the most common and lethal subtype of ovarian cancer, characterized by high recurrence rates and limited treatment options following chemotherapy resistance. Its significant heterogeneity poses major challenges for effective therapy and clinical outcomes.

In this study, we present a systems-level analysis of spatial transcriptomics data to characterize tumor heterogeneity in post-neoadjuvant chemotherapy HGSOC patients. By integrating gene expression profiles with spatial localization and histological context, we quantified hallmark pathway activities across tissue regions. The computed pathway scores were then used for clustering to investigate intra-tumoral heterogeneity. We also constructed gene co-expression network within tumor-enriched regions. Finally, we examined the association of these co-expressed modules with treatment response.

Clustering based on pathway activity scores revealed spatially distinct regions enriched for different hallmark pathways, uncovering functionally diverse cellular subpopulations within the tumor microenvironment. Tumor cell-enriched clusters show difference in pathways related to proliferation, metabolism, immune signaling and stress response, while fibroblast-rich regions exhibit upregulation of epithelial-mesenchymal transition (EMT). Unsupervised co-expression analysis further revealed gene modules associated with both biological processes and clinical phenotypes. Poor responders exhibit higher expression of gene modules involved in stress response, ribosomal function, oxidative phosphorylation, and cell-cycle regulation. In contrast, good responders show elevated activity in modules enriched for immune activation, extracellular matrix (ECM) remodeling, and inflammatory signaling.

Our findings provide insights into spatially resolved functional states, tumor heterogeneity, and molecular features associated with treatment response, offering a foundation for precision oncology approaches in ovarian cancer.

## Introduction

Ovarian cancer is one of the most aggressive and lethal gynecological malignancies. It encompasses a heterogeneous group of malignancies originating from various cell types, including epithelial, stromal, and germ cells, with epithelial ovarian cancer (EOC) accounting for approximately 95% of total cases^1^. Among EOCs, high-grade serous ovarian carcinoma (HGSOC) is the most prevalent and aggressive histologic subtype, comprising 70% of all EOC cases. These tumors are believed to originate more frequently from the fallopian tube rather than the ovarian surface epithelium and are often linked to serous tubal intraepithelial carcinoma as a precursor lesion^2^. According to findings from The Cancer Genome Atlas (TCGA) Project, approximately 10% of advanced HGSOC cases exhibit multiple regions of chromosomal gains or losses, leading to the amplification of over 30 growth-promoting genes^3^. Furthermore, mutations in the TP53 gene are hallmark characteristic of HGSOC, occurring in over 90% of cases. BRCA1/2, key players in homologous recombination (HR) repair, are inactivated in around 20% of patients through mechanisms such as germline or somatic mutations and epigenetic silencing^2^. Additionally, HGSOC is highly heterogeneous, with most alterations occurring in only a small subset of tumors^4,5^.

HGSOC is typically diagnosed at advanced stages (III or IV), when the cancer has already spread within or beyond the abdominal cavity^6^. The early detection of high-grade serous ovarian carcinoma (HGSOC) is challenging due to the absence of specific symptoms, which are often overlooked or misattributed to other conditions. One of the earliest biomarkers identified for HGSOC is CA125, a transmembrane cell-surface protein encoded by the MUC16 gene. Elevated CA125 levels are observed in HGSOC compared to healthy ovarian tissue, making it the most used molecular marker^7^. However, CA125 has several limitations: it is elevated in only 23-50% of early-stage cases and is not consistently detectable in advanced HGSOC^8^. The late diagnosis significantly reduces therapeutic efficacy and overall survival rates^9^. HGSOC is also associated with a poor prognosis, driven not only by late-stage presentation but also by the development of chemoresistance^10^. Most patients with advanced-stage HGSOC undergo cytoreductive surgery followed by platinum- and taxane-based chemotherapy. While initial response rates are high (∼80%), most patients eventually relapse due to the emergence of chemoresistance, severely limiting effective treatment options^10^. Therefore, characterizing intra-tumoral heterogeneity is crucial in HGSOC, as it is closely associated with recurrence, chemotherapy resistance, and poor prognosis.

Spatial transcriptomics (ST) has emerged as a transformative technology to map molecular changes within tissues. ST provides multimodal data that include histology, spatial location, and gene expression, enabling us to explore the complex interplay between cells and their microenvironment while preserving the structural integrity of the tissue. Recent studies have demonstrated the utility of ST in identifying transcriptional patterns, cell-cell interactions, and functional states within the tumor microenvironment (TME)^11^. For instance, Xie et al. highlighted the critical involvement of endothelial cells in mediating the transition from squamous to glandular histological subtypes in lung adenocarcinoma patients^12^. Few studies have analyzed ST data to characterize spatial diversity in HGSOC tumor tissue, providing insights into tumor heterogeneity and microenvironmental interactions^13,14^. Denisenko et al. revealed extensive subclonal heterogeneity based on copy number alterations in HGSOC and suggested that distinct ligand-receptor expression patterns within subclones may influence how HGSOC cells interact with their surrounding microenvironment^14^.

In this study, we leveraged ST data to investigate the spatial landscape of hallmark pathways in HGSOC patients, aiming to characterize tumor heterogeneity, and TME. Figure 1 outlines the analytical workflow of the study. Our findings offer insights into cancer biology of post-neoadjuvant chemotherapy (NACT) patients and contributes to the advancement of the targeted therapeutic strategies.

**Figure 1:**
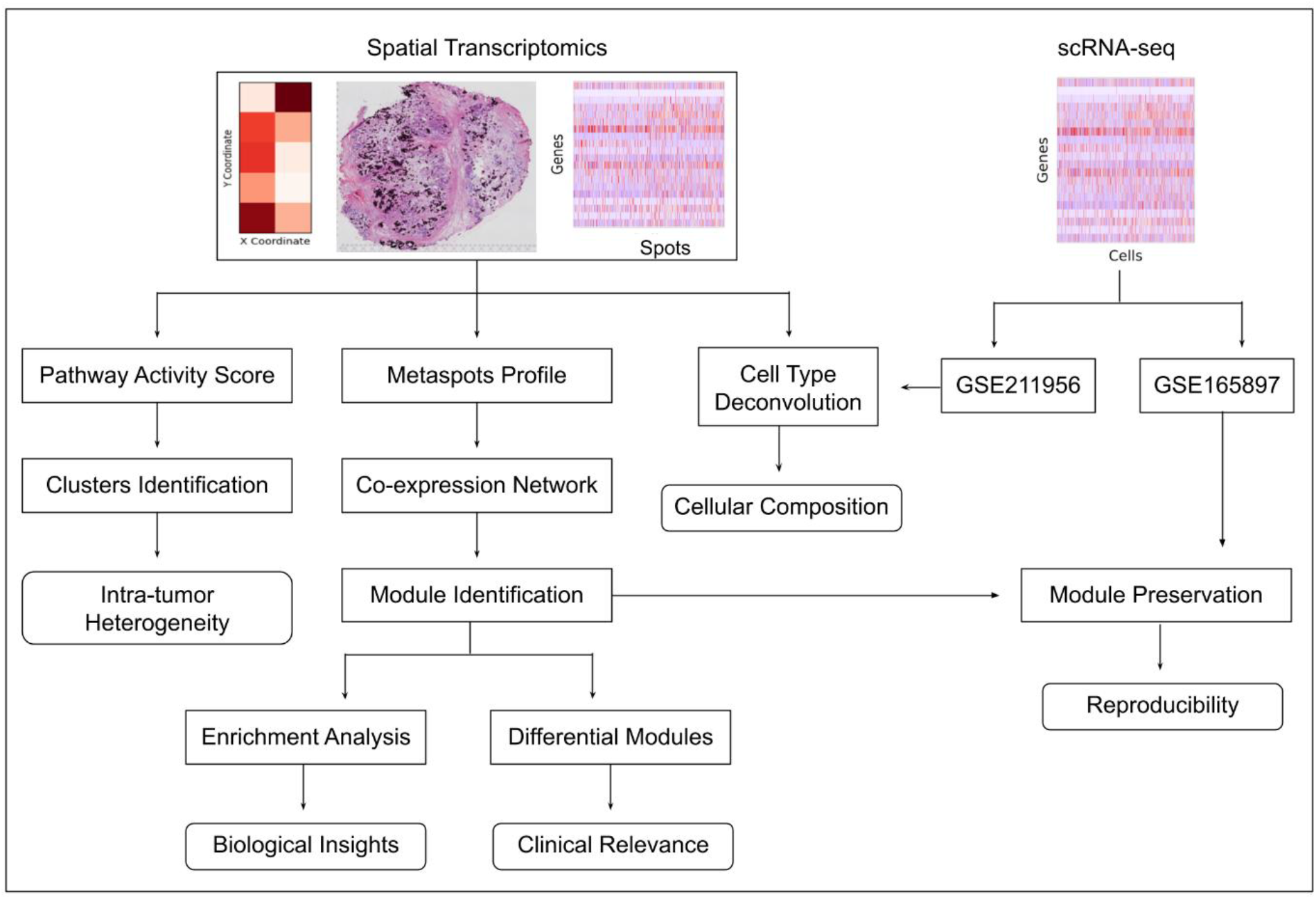
Overview of the ST-based workflow for characterizing intra-tumor heterogeneity and identifying functionally and clinically relevant gene modules in HGSOC. This workflow represents the stepwise approach used to interrogate ST data from cancer tissues. After inferring cellular composition, intra-tumor heterogeneity was explored by clustering spots based on pathway activity scores. Tumor-enriched spots were then used to identify co-expressed modules, followed by functional enrichment analysis. Module activity was also assessed in relation to chemotherapy response. Finally, preservation of modules was validated using single-cell RNA sequencing (scRNA-seq) data.

## Methods

### Dataset and Data Preprocessing

We downloaded 10x Genomics Visium ST datasets for ovarian cancer from the Gene Expression Omnibus (GEO) database^15^. The dataset (accession ID: GSE211956) includes samples from eight patients diagnosed with HGSOC, all of whom underwent 3-6 cycles of platinum-based chemotherapy^14^. Clinical data included chemotherapy response information based on the Chemotherapy Response Score (CRS), with patients classified as good (CRS 3), partial (CRS 2), or poor (CRS 1) responders. Specifically, three patients (P2, P3, P5) had a good response, three (P1, P7, P8) had a poor response, and two (P4, P6) had a partial response. Additionally, we utilized scRNA-seq data and corresponding cell type annotations from the original publication for cell type deconvolution. Detailed dataset information can be found in the original publication^14^.

For ST data, quality control was performed for each patient to ensure data reliability. Genes expressed in at least five spots were retained, and only spots with a minimum of 500 UMIs were included. Each spot was normalized by scaling its total counts across all genes, ensuring uniform total counts post-normalization.

We also downloaded another scRNA-seq data along with corresponding cell-type annotations, from treatment-naive and post-NACT paired samples of 11 homogeneously treated HGSOC patients (accession ID: GSE165897)^16^. The filtered barcodes obtained after quality control were available in the metadata file and were used in this study.

### Cell Type Deconvolution

Visium spots capture RNA from multiple cell types, and deconvolution helps determining the contributions of individual cell types. We performed cell type deconvolution using Robust Cell Type Decomposition (RCTD) to gain insights into the spatial arrangement and proportions of various cell types in the tissue. This method is one of the top-performing methods for cell type deconvolution^17^. RCTD utilizes reference scRNA-seq data to estimate the proportions of different cell types within each spatial spot based on their gene expression profiles^18^.

First, the scRNA-seq data were aggregated to create average gene expression profiles for each cell type, serving as reference templates that represent the transcriptional signatures of individual cell types. RCTD then models each spot as a combination of contributions from various cell types. A statistical model is applied to estimate gene expression for each cell type within a spot, assuming the observed gene counts follow Poisson distributions. This model is optimized using maximum likelihood estimation (MLE). RCTD also accounts for technical differences between sequencing platforms^18^. This method was implemented using the spacexr (v2.2.1) package available in R.

### Calculation of Pathway Activity Score

We adapted the STAN (Spatially informed Transcription factor Activity Network) framework to compute spot-specific pathway activity scores. STAN is originally designed to infer transcription factor (TF) activity by integrating ST data with known TF-target gene relationships to construct spatially aware regulatory networks^19^. To extend this framework for pathway-level analysis, we replaced the TF-target gene sets with curated pathway gene sets obtained from MSigDB^20^.

For each spatial location, the pathway activity scores were estimated based on the spatially weighted ridge regression problem. Let *Y* ∈ ℝ^*N*×*M*^ denote the normalized gene expression matrix, where *N* is the number of genes and *M* the number of spots, and let *D* ∈ ℝ^*N*×*p*^ represent a prior matrix linking genes to pathways, with *p* denoting the number of pathways. The core regression problem aims to estimate a pathway activity matrix by learning weight matrix *W* ∈ ℝ^*p*×*M*^ such that:

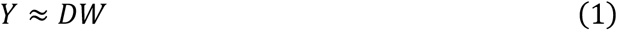

To incorporate spatial and morphological context, each spot *i* is represented by a five-dimensional feature vector *v*(*i*) = (*x*_*i*_, *y*_*i*_, *ωr*_*i*_, *ωg*_*i*_, *ωb*_*i*_), where (*x*_*i*_, *y*_*i*_) are spatial coordinates; (*r*_*i*_, *g*_*i*_, *b*_*i*_) are RGB values of local image patches centered on each spot and *ω* is weight parameter of the image features. All components are normalized to zero mean and unit variance across all spots. Next, spatial and morphological similarity between spots was encoded using a Gaussian kernel *K* ∈ ℝ^*M*×*M*^ as follows:

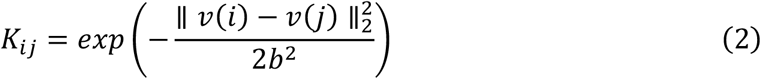

where *b* is the bandwidth parameter of the Gaussian kernel. Pathway activity is then estimated by augmenting the Equation 1 with regularization terms that separate spatially structured and unstructured components of the activity matrix. The spatially weighted ridge regression is formulated as follows:

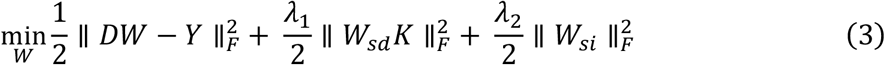

where *W* = *W*_*sd*_ + *W*_*si*_, with *W*_*si*_ capturing non-spatial variation and *W*_*sd*_ capturing spatially smoothed activity; *F* denotes the Frobenius norm; and *λ*_1_and *λ*_2_ are regularization parameters. The solution is obtained via low-rank approximation of *K*, as described in the original STAN paper^19^. For benchmarking, the same regression problem without spatial features was solved using ridge regression. The performance was assessed using 10-fold cross-validation by computing the Pearson correlation coefficient (PCC) between predicted and observed gene expression profiles at spatial locations. The implementation was carried out in python using the scanpy (v1.10.1) and squidpy (v1.5.0) libraries. The code components were adapted from the STAN GitHub repository, where applicable^19^.

### Identification of Clusters based on Hallmark Pathways

To explore spatial variation in pathway activity, z-scored pathway activity scores across spots were used as input for principal component analysis (PCA), retaining the top 10 components. A nearest neighbor graph was constructed based on the PCA representation, and Leiden clustering was applied to identify spatial domains. UMAP was used to embed pathway activity patterns in two dimensions. We also overlaid clustering results on spatial plots to visualize the spatial organization of pathway activities. All analyses were performed in python using the scanpy (v1.10.1) library.

### Co-expression Network Analysis

High-dimensional co-expression network analysis (hdWGCNA) was performed across tumor spots in the ST dataset using the R package hdWGCNA (v0.4.4)^21,22^. Prior to network construction, the ST data were preprocessed using Seurat’s (v5.1.0) standard pipeline, and batch effects were corrected using Harmony (v1.2.1) to meet the prerequisites for downstream analysis^23,24^. Genes expressed in at least 5% of spots were retained, resulting in a set of 11,729 genes for network construction. To address data sparsity, hdWGCNA aggregates neighboring spots based on spatial coordinates to form metaspots, and then normalizes the resulting metaspot expression matrix^21^. The soft-thresholding power was chosen by evaluating the scale-free topology model fit across a range of values, selecting the smallest power for which the fit exceeded 0.8. Co-expression network construction and module detection were performed using the ConstructNetwork function with default parameters. For each identified module, module eigengene expression was calculated to summarize the overall expression pattern. Hub genes were defined based on eigengene-based connectivity (kME).

We also performed differential module eigengene (DME) analysis using the Wilcoxon rank-sum test to identify modules differentially expressed between good and poor responders. Modules with an adjusted p-value < 0.05 (Benjamini-Hochberg correction) were considered statistically significant.

Finally, module preservation analysis was performed to evaluate the reproducibility of the identified modules in an independent scRNA-seq dataset (GSE165897). To assess module preservation between the reference (ST) and query (scRNA-seq) datasets, the Z_summary_ statistic was used: a Z_summary_ ≥ 10 indicates strong preservation, 2 < Z_summary_ < 10 suggests moderate preservation, and Z_summary_ < 2 indicates no evidence of preservation in the query dataset.

### Module Enrichment Analysis

To characterize the biological functions of each co-expressed module, enrichment analysis was performed using Enrichr (v 3.4) in R^25^. The background gene set consisted of all genes retained for network construction. We assessed enrichment against multiple gene set databases, including MSigDB Hallmark gene sets^20^, KEGG pathways^26^, Reactome pathways^27^, WikiPathways^28^, and Gene Ontology (GO) categories (Biological Process, Cellular Component, and Molecular Function)^29^. In addition, tumor marker gene sets from a published single-cell RNA-seq study were used to interpret modules^16^. Enrichment terms with an adjusted p-value < 0.05 were considered statistically significant.

## Results

### Inference of Cellular Composition

To investigate the spatial distribution and relative abundance of different cell types within the tissue, we applied RCTD for cell type deconvolution. We observed distinct patterns of cellular composition across patients, highlighting the diversity in tissue architecture (Figure 2). Patient P1 and P4 exhibits a predominance of tumor cells and fibroblasts. Patient P2 shows a more diverse cellular profile, characterized by the presence of tumor cells, fibroblasts, macrophages, and B/Plasma cells. Patient P3 is primarily composed of fibroblasts and tumor cells, with limited involvement of myofibroblasts. Patient P5 is notable for being predominantly composed of macrophages, with only a small proportion of tumor cells and fibroblasts, reflecting a unique cellular distribution compared to the other samples. In Patient P6, fibroblasts and macrophages were the dominant cell types, while tumor cells and B/plasma cells appeared in lesser abundance. Patient P7 is almost entirely composed of tumor cells, with minimal presence of fibroblasts. Patient P8 exhibits a more complex cellular composition, with contributions from tumor cells, fibroblasts, myofibroblasts, and endothelial cells. This analysis provides a comprehensive view of how different cell types are spatially organized within the tissue, shedding light on their arrangement within the microenvironment. Such insights are essential for uncovering patterns of cellular heterogeneity, tissue architecture, and the underlying biological processes unique to each sample post-treatment.

**Figure 2:**
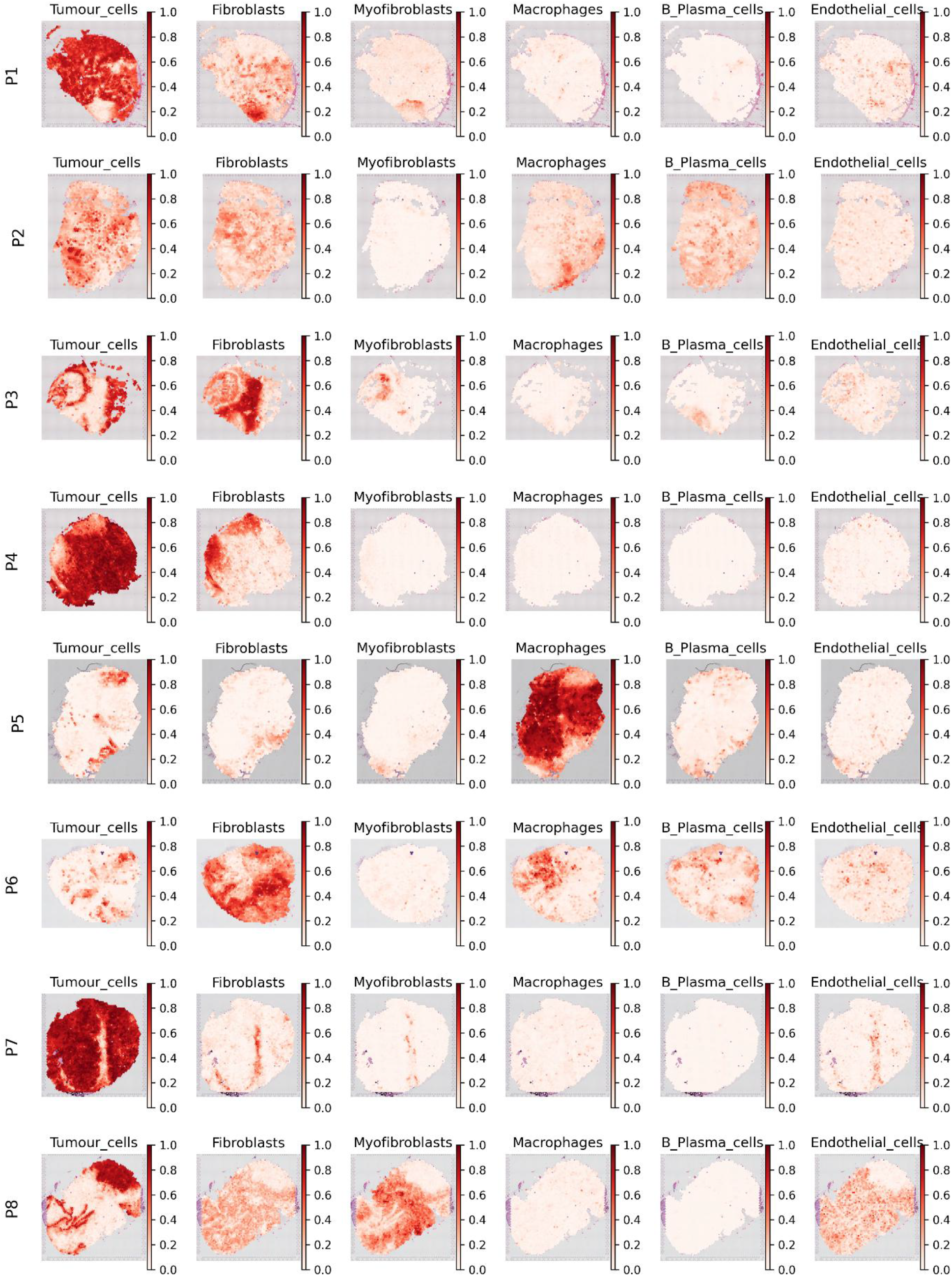
Mapping of the proportions of individual cell types across the spots in tissue sections from HGSOC samples. The colour bar indicates the weights assigned to each cell type, representing their relative contributions within each spot.

### Characterization of Intra-tumoral Heterogeneity Based on Pathway Activities

To identify different functional state within cancer tissue, we computed pathway activity scores by using ST data. Gene expression along with H&E image and spatial location were processed to generate activity scores for curated gene sets derived from the MSigDB Hallmark collection, enabling quantification of pathway activation at the sample level. The resulting pathway scores were subsequently utilized for clustering analysis to explore intra-tumoral heterogeneity. Our analysis reveals distinct regions sharing pathway activity patterns, which may correspond to biologically meaningful subpopulations within TME.

In patient P1, we identified four clusters based on the pathway activity profile (Figure 3). Clusters-1 and 2 corresponds to two distinct populations of tumor cells, while cluster-3 is primarily composed of fibroblasts. Cluster-0 contains a mix of tumor cells and fibroblasts. To gain deeper insights into the functional distinctions among clusters, we visualized the activity of hallmark pathways using a heatmap (Figure 3). In Cluster-0, we observed high activity of pathways associated with cell cycle, oxidative phosphorylation, DNA repair, survival signalling and enhanced biosynthesis. These pathways are often associated with therapy resistance. Cluster-1 exhibits downregulation of hallmark pathways, suggesting a region potentially sensitive to chemotherapy. Cluster-2 shows elevated activity in KRAS signaling, apoptosis, complement, and reactive oxygen species (ROS) pathways, reflecting stress response and immune activation features. These features may indicate a response to chemotherapy. In cluster-3, we noted upregulation of epithelial-mesenchymal transition (EMT), myogenesis, coagulation, and TGF-β signaling pathways. This fibroblast-rich cluster likely represents a reactive stromal component that contributes to tumor progression and immune evasion. It has been observed that TGF contributes to advanced HGSOC progression by modulating the activity of cancer-associated fibroblasts signatures in TME^30^. A detailed visualization of hallmark pathway activity across clusters in each sample is provided in Supplementary Figures (S1-S7).

**Figure 3:**
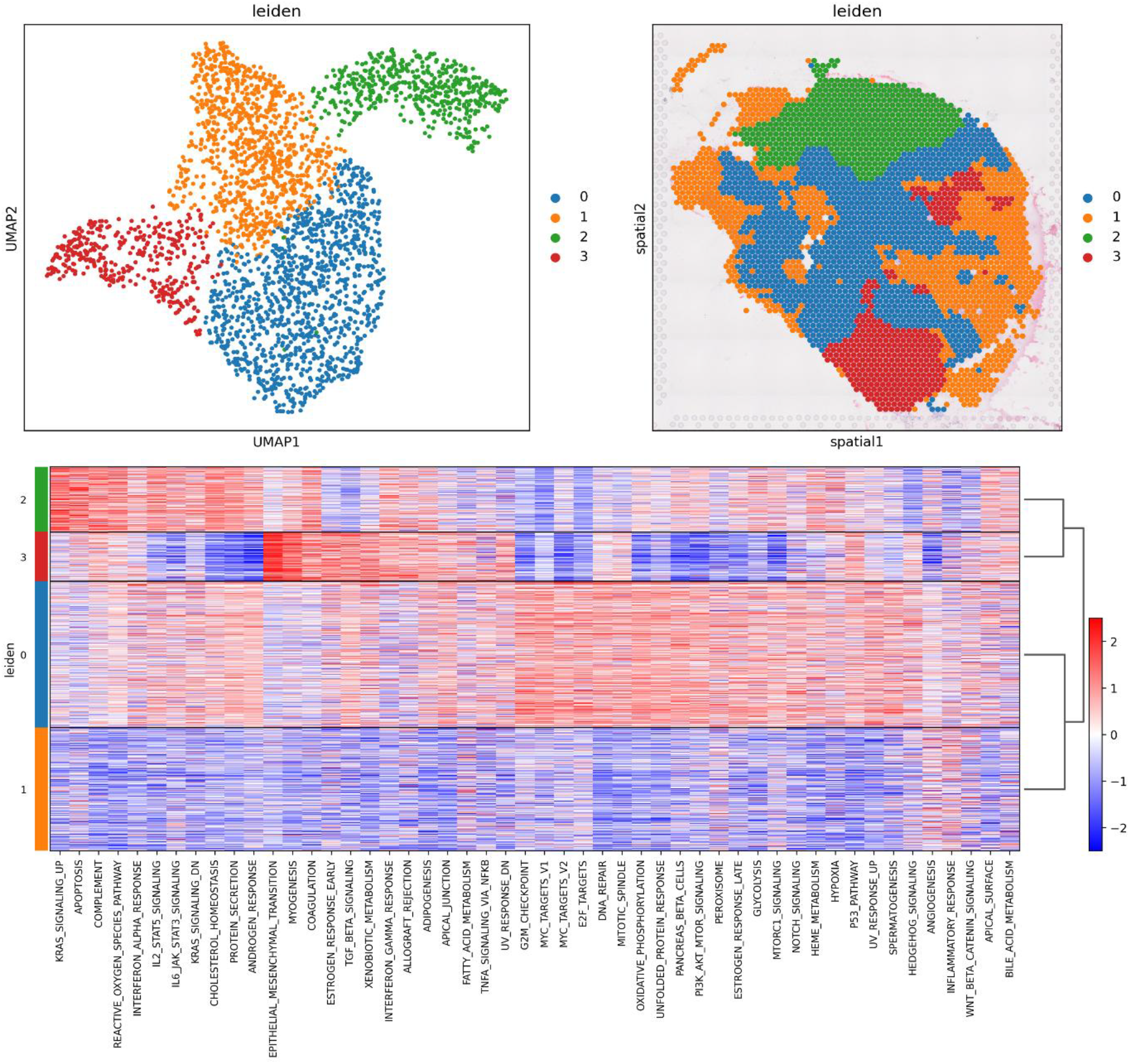
Clustering and pathway activity landscape in patient P1. UMAP and spatial plots display the clustering of spots, while the heatmap illustrates the activity scores of hallmark pathways across identified clusters.

In the seven remaining patients, our analysis identifies four clusters in five samples (P2, P3, P4, P7, and P8), and three clusters in two samples (P5, P6) (Figure S1-S7). Key observations across these patients are summarized below, with emphasis on distinct cellular compositions and hallmark pathway activity patterns within the identified clusters. In patients P2 and P5, we identified macrophage-enriched cluster-3 in P2 and clusters 0 and 1 in P5 (Figure S1 and S4). Cluster-3 in P2, and cluster-1 in P5, shows upregulation of pathways related to ROS, complement activation, and KRAS signaling. Additionally, cluster-1 of patient P5 exhibits high expression of cholesterol homeostasis, bile acid metabolism, and TNFA signaling via NFKB. In contrast, cluster-0 of P5 shows downregulation across all hallmark pathways.

In patient P3, clusters 0 and 3 were largely composed of fibroblasts (Figure S2). Cluster-0 displays elevated activity of EMT and coagulation pathways, whereas cluster-3 shows overall low pathway activity, which may reflect tissue remodeling following tumor clearance. Similarly, cluster-0 in patient P6, which was fibroblast-enriched, shows upregulation of EMT and coagulation (Figure S5). In patients P4 and P7, we observed that three clusters were composed predominantly of tumor cells, while one cluster comprised a mixture of tumor cells and fibroblasts (Figure S3 and S6). Specifically, in patient P7, cluster-0 exhibits negligible pathway activity, cluster-1 shows general downregulation, and cluster-2 demonstrates upregulation in most hallmark pathways. Cluster-3, which consists of both fibroblast and tumor-enriched spots, shows upregulation in pathways such as EMT, angiogenesis, myogenesis, coagulation, and hypoxia.

Overall, we observed elevated activity of pathways related to cell cycle, stress response, immune signaling, and cellular metabolism within tumor regions, whereas fibroblast-rich areas show higher activity of pathways such as EMT, myogenesis and coagulation. Spatial visualization of key pathway activity scores highlights the underlying heterogeneity within cancer tissues (Figure 4).

**Figure 4:**
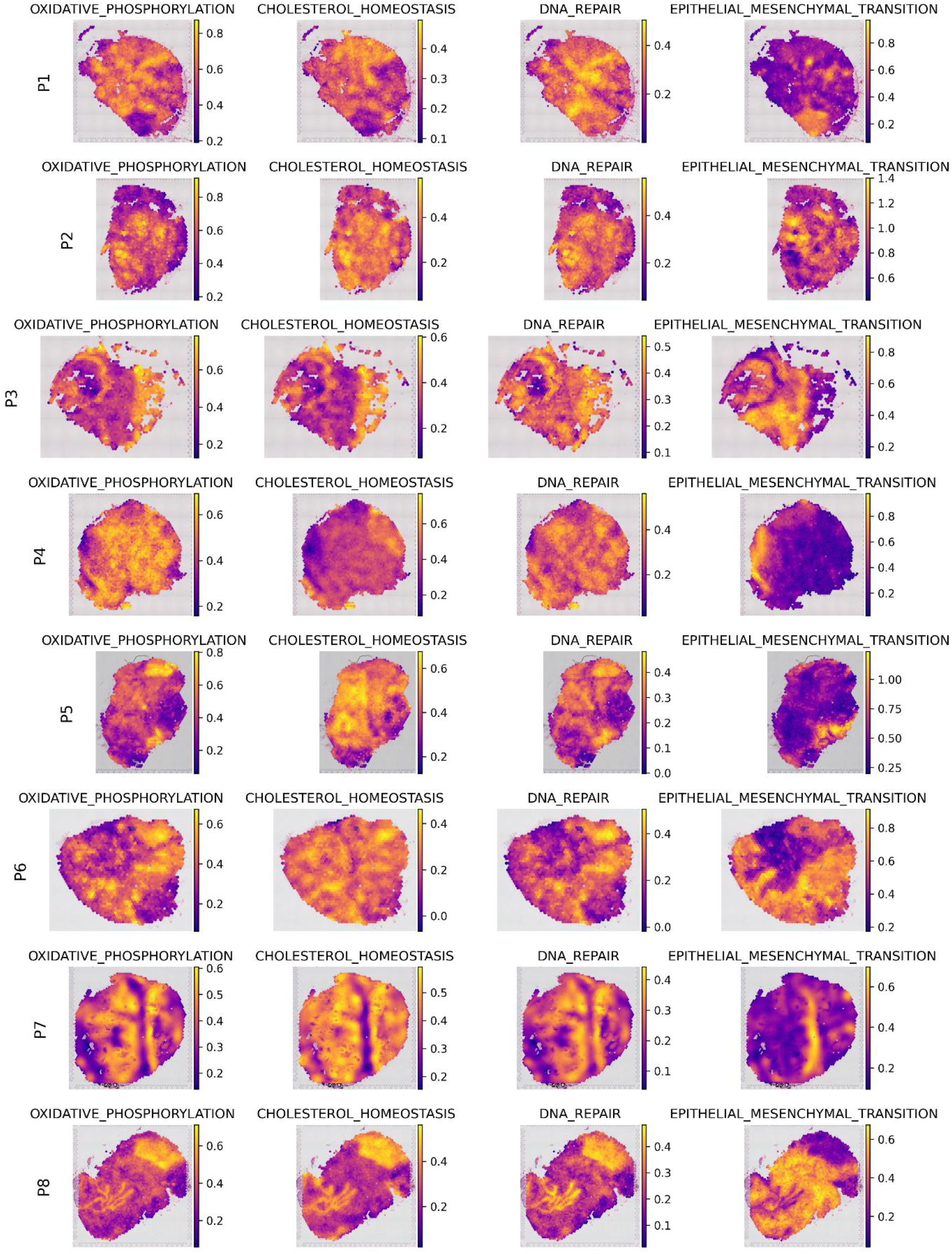
Spatial distribution of key hallmark pathway activity scores across different patients, highlighting intra-patient transcriptional heterogeneity. The color scale adjacent to each plot indicates the continuous range of pathway activity scores, with warmer colors denoting higher activity levels.

### Identification of Co-expressed Modules

In the previous analysis, we examined individual samples to investigate molecular alterations within TME. We next extended our analysis to include tumor spots from all samples in order to explore gene-gene expression pattern within tumor cells. A weighted gene co-expression network was constructed by integrating spatial location and gene expression data. Figure 5A illustrates the selection of soft-thresholding powers used to assess network topology. A soft power of 9 was selected for downstream network construction, resulting in the identification of 7 co-expression modules (Figure 5B). Each module is characterized by its eigengene expression, which represents the principal component of gene expression within the module (Table S1). The spatial distribution of eigengene values across tumor spots reflects the activity of these modules in tissue regions (Figure S8). Based on eigengene connectivity, hub genes were identified for each module. Figure 5C shows the co-expression network of the top hub genes from Module 1. The module includes hub genes associated with antigen presentation (e.g., HLA-A, HLA-B, HLA-C, HLA-DRA, HLA-DRB1, B2M), immunoglobulin production (IGHA1, IGHG1, IGHM, IGKC, IGLC2), and complement activation (C1QA, C1QB, C1QC), suggesting the presence of immune-related signaling within tumor spots. The networks for the remaining modules are provided in the supplementary material (Figure S9).

**Figure 5:**
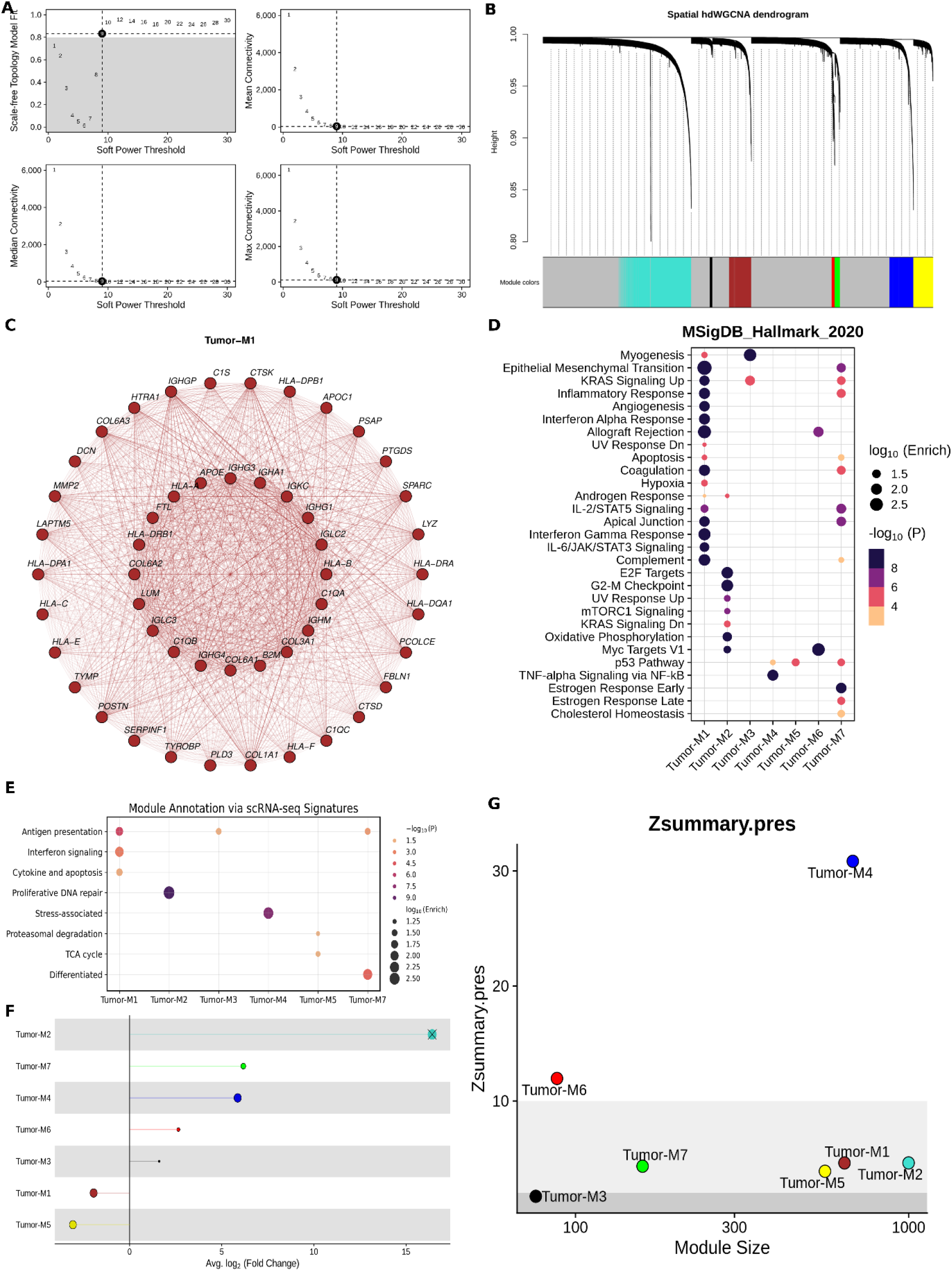
Co-expression network analysis of tumor spots in HGSOC. (A) Selection of soft-thresholding power based on scale-free network topology. (B) Dendrogram plot showing the identification of spatially co-expressed gene modules. (C) Network plot of the 50 hub genes in Module 1. (D) Dot plot showing module enrichment for MSigDB hallmark gene sets. (E) Dot plot showing module enrichment for tumor-specific gene signatures reported in a previous scRNA-seq study. (F) Lollipop plot displaying differentially expressed modules between good and poor responders. (G) Z-summary scores indicating preservation of identified modules in an independent scRNA-seq dataset.

To gain insight into the biological relevance of these modules, we performed enrichment analysis using multiple curated gene sets, including the MSigDB hallmark gene signatures (Figure 5D and Figure S10-15). Module 1 is functionally enriched for pathways related to EMT, immune signaling, and extracellular matrix (ECM) remodeling. Consistent patterns were observed across KEGG (antigen processing and presentation, ECM receptor interaction, cell adhesion molecules, allograft rejection), WikiPathways (type II interferon signaling, macrophage markers, allograft rejection), and Reactome (complement activation, ECM organization) (Figure S10-12). Gene Ontology (GO) analysis further highlighted enrichment in MHC class II-related protein complexes, chemokine activity, and ECM structural components (Figure S13-15). Notably, we observed that the majority of EMT-enriched genes in this module are EMT activators and include transcription factors (SNAI2, PRRX1), signaling molecules (TGFB1), and matrix remodeling enzymes.

Module 2 is predominantly enriched in cell cycle-related processes, oxidative phosphorylation, mTORC1 signaling and GO terms such as mitotic spindle assembly checkpoint, mitochondrial electron transport chain, and mitochondrial inner membrane. Module 3 shows enrichment in myogenesis, and collagen-containing ECM. Module 4 is enriched in pathways such as TNF-alpha signaling via NF-κB, oxidative stress response, and cilium assembly. Module 6 is enriched for ribosome-related processes, translation, and macromolecule biosynthesis. Enriched GO terms highlighted ribosomal subunits, rRNA binding, and components of the rough ER, suggesting an active role in protein synthesis machinery. Module 7 shows enrichment in estrogen response (early and late).

A recent scRNA-seq study of chemotherapy-treated patients reported tumor-specific gene signatures. To evaluate the biological relevance and overlap of our spatial co-expression modules with those tumor signatures, we performed enrichment analysis using their reported gene sets. Our analysis reveals consistency between the two datasets (Figure 5E). Specifically, module 1 is significantly enriched for antigen presentation, interferon signaling, and cytokine and apoptosis-related signatures. Module 2 is enriched for proliferative DNA repair programs. Module 4 is associated with stress-related signatures. Module 5 is enriched in proteasomal degradation and TCA cycle processes. Finally, module 7 is enriched for differentiation-associated programs.

We next investigated the relationship between the co-expressed gene modules and chemotherapy response. Six modules are found to be differentially expressed between good and poor responders (Figure 5F). Specifically, modules 3, 4, 6, and 7 are upregulated in poor responders, whereas modules 1 and 5 shows higher expression in good responders. Although module 2 is not statistically significant, it exhibits a high log fold change in poor responders.

To assess the robustness and reproducibility of the identified co-expression modules, we conducted module preservation analysis using the scRNA-seq dataset employed earlier for gene signature enrichment. Modules 4 and 6 shows strong preservation, indicating high consistency across datasets. Modules 1, 2, 5, and 7 exhibits moderate preservation, suggesting partial concordance in gene co-expression pattern. In contrast, module 3 is poorly preserved, implying it may represent a dataset-specific or context-dependent signal (Figure 5G).

## Discussion

Although treatment approaches for advanced HGSOC have progressed, clinical outcomes remain poor. Early relapse, therapy resistance, and a shortage of effective therapeutic targets continue to be major challenges in improving overall survival rates. Recent advances in scRNA-seq and ST technologies have enabled deeper insight into TME and its spatial organization, particularly in the context of chemotherapy^13,14,16,31^. Single-cell analyses of HGSOC reported extensive intratumor heterogeneity in epithelial cancer cells following neoadjuvant chemotherapy^31^. Another scRNA-seq study identified stress-associated transcriptional programs contributing to chemoresistance in metastatic HGSOC^16^. Leveraging the ST dataset, our study investigates the pathway landscape from the perspective of spatially-resolved pathway activity and gene co-expression pattern, revealing biologically and clinically relevant changes in the HGSOC (Figure 1).

To systematically investigate the pathway landscape in HGSOC, we first characterized intra-tumor heterogeneity through spatial variation in hallmark pathway activity within cancer tissues. By integrating gene expression profiles with spatial coordinates and histological context, we quantified pathway activity at each spot. This approach allowed us to move beyond individual gene-level variation and capture coordinated changes in key hallmark processes. It also facilitated the identification of distinct pathways activated in different regions, revealing their associations with specific cell types within TME. Our findings highlight spatial heterogeneity in pathway activity within cancer tissue (Figure 2-3 and S1-S7). Clustering of pathway activity reveals that tumor cell-enriched clusters generally exhibit high activity of hallmark processes such as cell cycle regulation, DNA repair, stress response, immune signalling, and oxidative phosphorylation, while fibroblast-rich regions consistently show increased EMT, coagulation, and myogenesis (Figure 4). Multiple studies have demonstrated that activation of cellular stress pathways, such as UPR, promotes chemoresistance by increasing the expression of antioxidant genes, efflux pump genes, and DNA damage response genes^32,33^. Upregulated antioxidants detoxify ROS produced by chemotherapeutic agents, allowing tumor cells to survive oxidative stress^32^. Our findings also support prior evidence that cancer-associated fibroblasts are major drivers of EMT within TME^34^. However, this pattern was observed in both good and poor responders. In good responders (e.g., patient P3), this may reflect tissue remodeling following tumor clearance, whereas in poor responders (e.g., patients 1 and 7), they are more likely associated with invasion and therapy resistance. Variation in activity of oxidative phosphorylation and cholesterol homeostasis aligns with our findings from previous single cell metabolic study of HGSOC^35^ (Figure 4). Additionally, macrophage-enriched clusters, as observed in patients P2 and P5, display activation of inflammatory and stress response pathways (Figure S1 and S4). These spatial differences reflect functional compartmentalization within TME, influenced by both malignant and stromal components.

We further extended our analysis to tumor spots across the samples to capture coordinated expression pattern within the tumor cells. Using co-expression network analysis within tumor-enriched spots, we identified gene modules that reflect disease-wide expression patterns in HGSOC (Figure 5A-C). These modules are enriched for biological processes including proliferation, stress response, metabolism, differentiation, EMT, and immune-related pathways (Figure 5D-E). These observations align with our first part of pathway analysis. We also observed these modules to be significantly differentially expressed between good and poor responders, suggesting their clinical relevance (Figure 5F). Notably, modules associated with stress pathways, myogenesis, ribosome-related processes, oxidative phosphorylation, and cell-cycle pathways are linked to poor responders, while those enriched for ECM remodeling, immune, and inflammatory signaling are more expressed in good responders. Evidence from multiple studies suggests that ribosome biogenesis-related proteins contribute to chemoresistance in cancer^36^. For instance, knockdown of RPL13 has been shown to suppress cell proliferation, while its overexpression promotes resistance to chemotherapy in gastric cancer^37^. RPL13 was identified as one of the top hub genes in our co-expression module M6. Oxidative phosphorylation has also been shown to be essential for the development of chemoresistance across multiple tumor types, and targeting this metabolic pathway can selectively eliminate cancer stem cells and delay resistance acquisition^38^. In triple-negative breast cancer (TNBC), resistant cells rely on oxidative phosphorylation for energy production compared to chemotherapy-sensitive cells^39^. On the other hand, several studies have highlighted the favourable role of immune signaling in promoting good responses to chemotherapy^40,41^. Chemotherapy has been shown to induce immunogenic cell death, a process that enhances antitumor immune responses by promoting the release of damage-associated molecular patterns (DAMPs)^42^.

Finally, the preservation of modules in independent scRNA-seq data demonstrates the reproducibility of our findings (Figure 5G). Enrichment of these modules with gene signatures derived from scRNA-seq study further validated their biological significance and confirms the finding of tumor-specific stress-associated transcriptional programs in HGSOC^16^. These results suggest that the co-expression patterns captured in the ST data reflect biologically meaningful programs.

## Conclusion

Our study highlights a systems-level analysis of ST data to uncover the spatial landscape of hallmark pathways in HGSOC. Pathway-level analysis reveals intra-tumor heterogeneity and identifies spatially distinct functional states within TME, while co-expression network analysis uncovers co-expressed modules with biological and clinical relevance. These complementary approaches capture distinct yet overlapping aspects of tumor biology, shedding light on functional states and their spatial organization within cancer tissue. These findings provide deeper insight into tumor heterogeneity in post-NACT ovarian cancer patients and contribute to advancing precision therapeutic approaches.

## Data Availability

All datasets used in this study are available on NCBI GEO: ST (GSE211956) and scRNA-seq (GSE165897).

## Code Availability

The packages used in the analysis are referenced in the manuscript. The source codes is available upon request.

## Author Contributions

Conceptualization: PKV; Methodology: AS; Formal analysis and investigation: AS; Writing - original draft preparation: AS; Writing - review and editing: AS, PKV; Funding acquisition: P KV; Supervision: PKV.

## Acknowledgements

This work was supported by iHUB-Data, International Institute of Information Technology, Hyderabad, India. The funding body has no role in the study design and analysis.

## Competing interests

The authors declare that they have no competing interests.

## Notes

### Competing Interest Statement

The authors have declared no competing interest.

## Reference

1. Lheureux, S., Braunstein, M. & Oza, A. M. Epithelial ovarian cancer: Evolution of management in the era of precision medicine. CA Cancer J Clin 69, 280–304 (2019).

2. Krzystyniak, J., Ceppi, L., Dizon, D. S. & Birrer, M. J. Epithelial ovarian cancer: the molecular genetics of epithelial ovarian cancer. Annals of Oncology 27, i4–i10 (2016).

3. Bell, D. et al. Integrated genomic analyses of ovarian carcinoma. Nature 2011 474:7353 474, 609–615 (2011).

4. Azzalini, E., Stanta, G., Canzonieri, V. & Bonin, S. Overview of Tumor Heterogeneity in High-Grade Serous Ovarian Cancers. International Journal of Molecular Sciences 2023, Vol. 24, Page 15077 24, 15077 (2023).

5. Veneziani, A. C. et al. Heterogeneity and treatment landscape of ovarian carcinoma. Nature Reviews Clinical Oncology 2023 20:12 20, 820–842 (2023).

6. Berek, J. S., Renz, M., Kehoe, S., Kumar, L. & Friedlander, M. Cancer of the ovary, fallopian tube, and peritoneum: 2021 update. International Journal of Gynecology & Obstetrics 155, 61–85 (2021).

7. Köbel, M. et al. Ovarian Carcinoma Subtypes Are Different Diseases: Implications for Biomarker Studies. PLoS Med 5, e232 (2008).

8. Dochez, V. et al. Biomarkers and algorithms for diagnosis of ovarian cancer: CA125, HE4, RMI and ROMA, a review. J Ovarian Res 12, 1–9 (2019).

9. Bowtell, D. D. et al. Rethinking ovarian cancer II: reducing mortality from high-grade serous ovarian cancer. Nature Reviews Cancer 2015 15:11 15, 668–679 (2015).

10. Wiedemeyer, W. R., Beach, J. A. & Karlan, B. Y. Reversing platinum resistance in high-grade serous ovarian carcinoma: Targeting BRCA and the homologous recombination system. Front Oncol 4 MAR, 80581 (2014).

11. Jin, Y. et al. Advances in spatial transcriptomics and its applications in cancer research. Molecular Cancer 2024 23:1 23, 1–24 (2024).

12. Xie, L. et al. Spatial transcriptomics reveals heterogeneity of histological subtypes between lepidic and acinar lung adenocarcinoma. Clin Transl Med 14, e1573 (2024).

13. Stur, E. et al. Spatially resolved transcriptomics of high-grade serous ovarian carcinoma. iScience 25, 103923 (2022).

14. Denisenko, E. et al. Spatial transcriptomics reveals discrete tumour microenvironments and autocrine loops within ovarian cancer subclones. Nature Communications 2024 15:1 15, 1–15 (2024).

15. Edgar, R., Domrachev, M. & Lash, A. E. Gene Expression Omnibus: NCBI gene expression and hybridization array data repository. Nucleic Acids Res 30, 207–210 (2002).

16. Zhang, K. et al. Longitudinal single-cell RNA-seq analysis reveals stress-promoted chemoresistance in metastatic ovarian cancer. Sci Adv 8, 1831 (2022).

17. Li, B. et al. Benchmarking spatial and single-cell transcriptomics integration methods for transcript distribution prediction and cell type deconvolution. Nature Methods 2022 19:6 19, 662–670 (2022).

18. Cable, D. M. et al. Robust decomposition of cell type mixtures in spatial transcriptomics. Nature Biotechnology 2021 40:4 40, 517–526 (2021).

19. Zhang, L. et al. STAN, a computational framework for inferring spatially informed transcription factor activity. bioRxiv 2024.06.26.600782 (2024) doi:10.1101/2024.06.26.600782.

20. Liberzon, A. et al. The Molecular Signatures Database Hallmark Gene Set Collection. Cell Syst 1, 417–425 (2015).

21. Morabito, S., Reese, F., Rahimzadeh, N., Miyoshi, E. & Swarup, V. hdWGCNA identifies co-expression networks in high-dimensional transcriptomics data. Cell Reports Methods 3, (2023).

22. Langfelder, P. & Horvath, S. WGCNA: An R package for weighted correlation network analysis. BMC Bioinformatics 9, 1–13 (2008).

23. Hao, Y. et al. Integrated analysis of multimodal single-cell data. Cell 184, 3573–3587.e29 (2021).

24. Korsunsky, I. et al. Fast, sensitive and accurate integration of single-cell data with Harmony. Nat Methods 16, 1289–1296 (2019).

25. Kuleshov, M. V. et al. Enrichr: a comprehensive gene set enrichment analysis web server 2016 update. Nucleic Acids Res 44, W90–W97 (2016).

26. Kanehisa, M., Furumichi, M., Sato, Y., Matsuura, Y. & Ishiguro-Watanabe, M. KEGG: biological systems database as a model of the real world. Nucleic Acids Res 53, D672–D677 (2025).

27. Milacic, M. et al. The Reactome Pathway Knowledgebase 2024. Nucleic Acids Res 52, D672– D678 (2024).

28. Agrawal, A. et al. WikiPathways 2024: next generation pathway database. Nucleic Acids Res 52, D679–D689 (2024).

29. Consortium, T. G. O. et al. The Gene Ontology knowledgebase in 2023. Genetics 224, (2023).

30. Yeung, T. L. et al. TGF-β Modulates ovarian cancer invasion by upregulating CAF-Derived versican in the tumor microenvironment. Cancer Res 73, 5016–5028 (2013).

31. Shen, Y. et al. The impact of neoadjuvant chemotherapy on the tumor microenvironment in advanced high-grade serous carcinoma. Oncogenesis 11, 1–12 (2022).

32. Madden, E., Logue, S. E., Healy, S. J., Manie, S. & Samali, A. The role of the unfolded protein response in cancer progression: From oncogenesis to chemoresistance. Biol Cell 111, 1–17 (2019).

33. He, J., Zhou, Y. & Sun, L. Emerging mechanisms of the unfolded protein response in therapeutic resistance: from chemotherapy to Immunotherapy. Cell Communication and Signaling 2024 22:1 22, 1–22 (2024).

34. Szabo, P. M. et al. Cancer-associated fibroblasts are the main contributors to epithelial-to-mesenchymal signatures in the tumor microenvironment. Scientific Reports 2023 13:1 13, 1– 13 (2023).

35. Srivastava, A. & Vinod, P. K. A single-cell network approach to decode metabolic regulation in gynecologic and breast cancers. NPJ Syst Biol Appl 11, 1–15 (2025).

36. Elhamamsy, A. R., Metge, B. J., Alsheikh, H. A., Shevde, L. A. & Samant, R. S. Ribosome Biogenesis: A Central Player in Cancer Metastasis and Therapeutic Resistance. Cancer Res 82, 2344–2353 (2022).

37. Kobayashi, T. et al. Activation of the ribosomal protein L13 gene in human gastrointestinal cancer. Int J Mol Med 18, 161–170 (2006).

38. Zhao, Z., Mei, Y., Wang, Z. & He, W. The Effect of Oxidative Phosphorylation on Cancer Drug Resistance. Cancers 2023, Vol. 15, Page 62 15, 62 (2022).

39. Uslu, C., Kapan, E. & Lyakhovich, A. OXPHOS inhibition overcomes chemoresistance in triple negative breast cancer. Redox Biol 83, 103637 (2025).

40. Wong, D. Y. Q., Ong, W. W. F. & Ang, W. H. Induction of Immunogenic Cell Death by Chemotherapeutic Platinum Complexes. Angewandte Chemie - International Edition 54, 6483–6487 (2015).

41. Wang, Y. J., Fletcher, R., Yu, J. & Zhang, L. Immunogenic effects of chemotherapy-induced tumor cell death. Genes Dis 5, 194–203 (2018).

42. Fucikova, J. et al. Detection of immunogenic cell death and its relevance for cancer therapy. Cell Death & Disease 2020 11:11 11, 1–13 (2020).

